# Hierarchical Modular Structure of the Drosophila Connectome

**DOI:** 10.1101/2022.11.23.517722

**Authors:** Alexander B. Kunin, Jiahao Guo, Kevin E. Bassler, Xaq Pitkow, Krešimir Josić

## Abstract

The structure of neural circuitry plays a crucial role in brain function. Previous studies of brain organization generally had to trade off between coarse descriptions at a large scale and fine descriptions on a small scale. Researchers have now reconstructed tens to hundreds of thousands of neurons at synaptic resolution, enabling investigations into the interplay between global, modular organization, and cell type-specific wiring. Analyzing data of this scale, however, presents unique challenges. To address this problem we applied novel community detection methods to analyze the synapse-level reconstruction of an adult fruit fly brain containing over 20 thousand neurons and 10 million synapses. Using a machine-learning algorithm, we find the most densely connected communities of neurons by maximizing a generalized modularity density measure. We resolve the community structure at a range of scales, from large (on the order of thousands of neurons) to small (on the order of tens of neurons). We find that the network is organized hierarchically and larger-scale communities are composed of smaller-scale structures. Our methods identify well-known features of the fly brain, including its sensory pathways. Moreover, focusing on specific brain regions, we are able to identify subnetworks with distinct connectivity types. For example, manual efforts have identified layered structures in the fan-shaped body. Our methods not only automatically recover this layered structure, but also resolve finer connectivity patterns to downstream and upstream areas. We also find a novel modular organization of the superior neuropil, with distinct clusters of upstream and downstream brain regions dividing the neuropil into several pathways. These methods show that the fine-scale, local network reconstruction made possible by modern experimental methods are sufficiently detailed to identify the organization of the brain across scales, and enable novel predictions about the structure and function of its parts.

## Introduction

Understanding how brains function requires understanding how they are wired, and how this wiring underpins neural computation [4]. Advances in biology, imaging, and machine learning have led to a proliferation of vast, highly detailed connectomes of brain tissue from insects [45], mammals [12], and humans [47]. Dense reconstructions of neural tissue at synaptic resolution present new opportunities to interrogate the wiring principles of different brains. However, the enormous volume of the data presents fundamental challenges since it is difficult to identify relevant features of a large network and understand how these features interact.

The brains of sufficiently complex animals, including insects, are composed of interacting structural units that often have distinct functions. These “communities” are sometimes, but not always, anatomically distinct and have traditionally been identified using painstaking methods that involve the tracing of individual cells and their connections. However, as the number and size of connectomes grows, we will need automated methods to uncover a brain’s functional units and their interaction. Networks of the size and connectedness of connectomes pose fundamental technical challenges to many existing network community detection methods, the speed and accuracy of which can scale poorly with network size and often cannot resolve structures below a certain limiting size [17].

Insect brains are an excellent target for methods that automatically identify the salient features of complex networks. They are small enough and stereotyped enough that complete, validated connectomes are within reach [45]. Much is known about the organization of insect brains already, as anatomists have been able to characterize them at the level of single cells, circuits, and regions, and understand the computational roles and interactions between these components [21, 31, 30]. Understanding the structure of insect brains also provides insights into general organizational and computational principles of other brains [20, 29, 48]. Recently, automated methods have accelerated and amplified the abilities of scientists to identify neurons and their interactions at high resolution and in large quantities [51]. The fly Hemibrain [45], a synapse-level reconstruction of roughly two thirds of the volume of the brain of an adult female *Drosophila* fruit fly, is the largest (by number of neurons) connectome published to date.

Here we use an automated method to uncover cell communities that may constitute different signaling paths, or may be devoted to distinct computations in the Hemibrain network. We identify communities of neurons as the sets of neurons that are more densely-connected than expected in a random network. The community structure is found by partitioning the neurons in a way that maximizes a modularity density measure [7, 8, 2, 9, 19]. To find the maximizing partition we employ a recently introduced machine-learning algorithmic scheme RenEEL [18] that enables fast and accurate analysis of the Hemibrain network. By increasing a tuneable parameter in our modularity density measure, we resolve the structure of the network on increasingly smaller scales.

We identify communities that range in size from thousands to only a few neurons and find a roughly hierarchical organization of the structure [24]. At the coarsest scale, we automatically identify well-known brain regions and functional networks. At smaller scales, our analysis reveals how these brain regions are organized into sub-networks. It also identifies specific connectivity patterns among modules and brain regions. For example, we automatically recover the layered structure of the fan-shaped body [21] in an unsupervised way. By considering the cell type composition of the communities, we are able to identify potentially biologically relevant networks and cell-type-specific wiring patterns. Thus, we provide a scalable, automated method to identify structure in connectomes composed of tens of thousands of neurons, or more.

## Results

To infer the community organization of a connectome, we treat it as an undirected graph whose nodes are neurons and whose edge weights are defined as the total number of synapses between neurons. By initially treating the connectome as undirected, we obtain a liberal measure of communities, and we subsequently evaluate directed motifs on this undirected scaffold. We discover these communities based on the strength and density of connections between individual cells. Strong connections between neurons may be important for information flow in the network, and highly-connected groups of neurons may represent distinct computational circuits. We therefore assume that groups of strongly interacting neurons are likely to form functional units and use a multi-resolution community detection method to identify groups of densely connected cells. To validate our clustering approach, we compare our results to previously identified structures in the fly connectome, and apply our method to the connectome of the mushroom body of a *Drosophila* larva.

Many biological networks are hierarchical. Our method is designed to uncover such organization in an unsupervised way, without assuming *a priori* that a network is structured hierarchically. We identify communities in the network by partitioning the network into clusters in a way that maximizes a global measure, the Generalized Modularity Density *Q_g_*(*χ*) [19]. This measure increases with the density of connections within clusters, and depends on a tuneable parameter *χ* ≥ 0 which governs the resolution scale of the communities identified. At *χ* = 0, *Q_g_* (0) is equivalent to the classical Modularity, *Q* [36], and a relatively small number of large communities are identified (eight in the Hemibrain). As *χ* increases, maximizing *Q_g_*(*χ*) identifies progressively smaller, more densely connected communities (Table 1, see Methods). Thus, the resolution scale of the community structure within the network varies with *χ*. The number of communities identified at any particular value of *χ* is not predetermined, but is a result of the optimization. The communities we identify at larger *χ* are generally subsets of the communities we identify at smaller *χ* and, thus, the community structure is generally hierarchical. To find the partition that maximizes *Q_g_*(*χ*), we use Reduced network Extremal Ensemble Learning (RenEEL) [18], a recently introduced machine-learning algorithmic scheme for graph partitioning that enables fast and accurate results for networks of the size and density of the Hemibrain [18]. At this scale, without this speed and accuracy of RenEEL, this study would not have been possible.

**Table 1.** Statistics of communities in the Hemibrain network found by maximizing Generalized Modularity Density, *Q_g_*(*χ*). Increasing *χ* results in smaller, more tightly-connected communities. Weighed density is the sum of edge weights divided by the number of possible edges within a community; mean and inner quartile range exclude communities consisting of a single node.

| $\chi$ | # communities | Mean Size | Mean Weighted Density (IQR) |
| --- | --- | --- | --- |
| 0.0 | 8 | 2716.6 | 0.60 (0.30 - 0.64) |
| 0.05 | 68 | 699.5 | 2.15 (0.63 - 1.60) |
| 0.1 | 175 | 277.4 | 2.70 (0.72 - 2.99) |
| 0.25 | 440 | 66.9 | 8.16 (1.00 - 5.00) |
| 0.5 | 1485 | 16.1 | 23.52 (2.00 - 12.33) |
| 0.75 | 3076 | 7.5 | 27.62 (3.48 - 20.00) |
| 1.0 | 4421 | 5.1 | 30.35 (4.33 - 23.00) |

### Modular Structure in the Larval Mushroom Body is Driven by Cell Type and Anatomy

To validate our method, we applied it first to the connectome of the mushroom body of a larval fruit fly [1]. This network consists of 365 neurons and is composed of two symmetric hemispheres with lobes extending in the anterior and dorsal direction (Figure 1A). At the coarsest level, *χ* = 0, our method partitioned the mushroom body into the left and right hemispheres, as well as a bilateral dorsal-posterior bundle of 34 neurons with projections to both halves (Figure 1A). In addition, it identified two small clusters, one in the anterior bridge connecting the two hemispheres and one in the left hemisphere. The larger cluster bridging the two halves consists of sensory and other projection neurons that often come in left/right pairs. The small cluster in the anterior bridge consists of a left-right pair of dopaminergic neurons and one output neuron, with strong internal connections and strong connections to both hemispheres. The five-neuron cluster consists of very young Kenyon cells in the left hemisphere with few synaptic connections to each other and to the remaining neurons.

**Figure 1.**
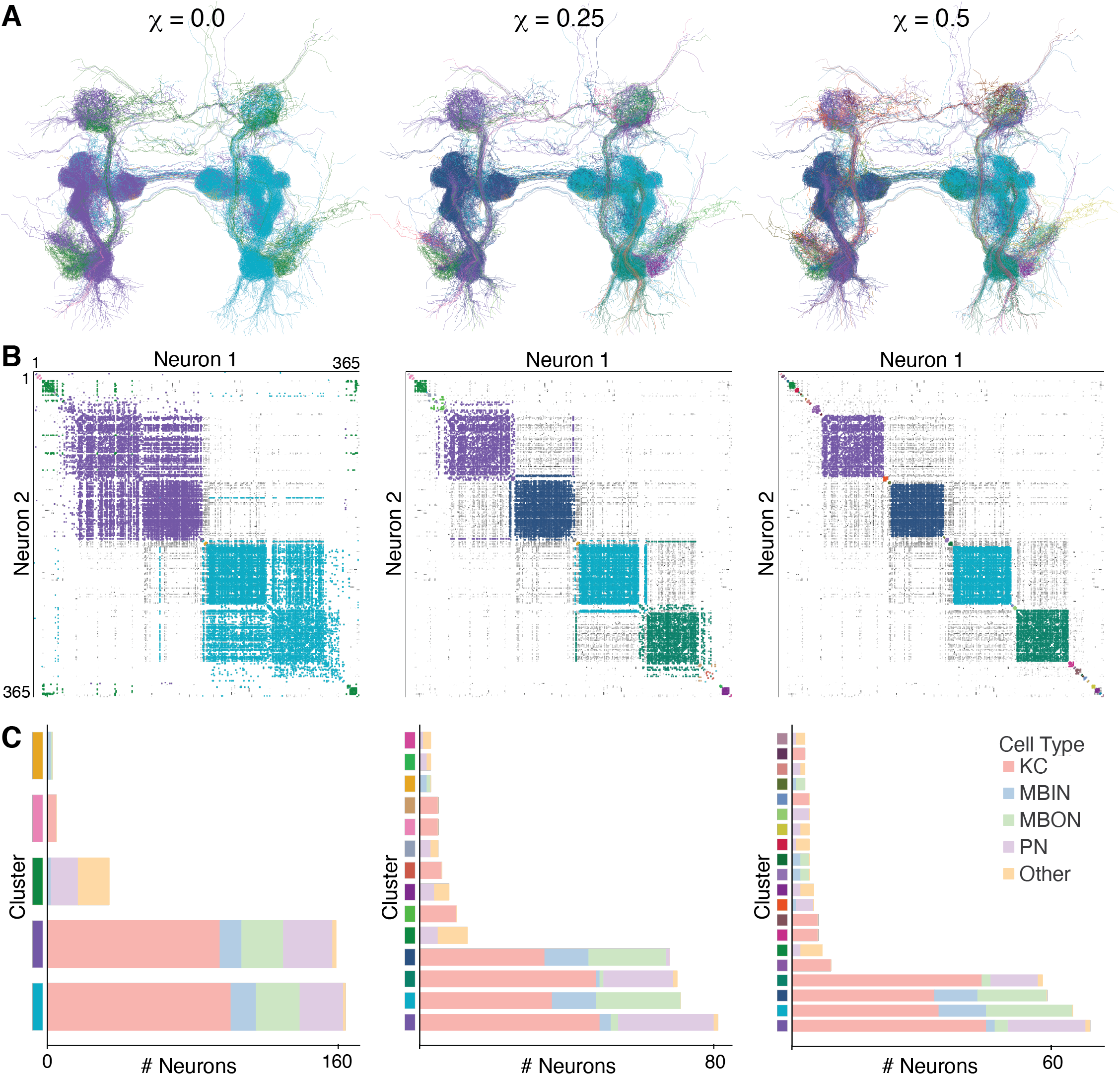
The larval mushroom body network exhibits hierarchical modularity which aligns with anatomy and cell type. **A.** Morphological reconstructions of the neurons in the larval mushroom body network, colored by cluster membership, viewed from a posterior and slightly dorsal viewpoint. From left to right, clustering was performed by maximizing generalized modularity density, *Q_g_*(*χ*), with *χ* = 0.0, 0.25, 0.5 (see Methods). **B.** Undirected adjacency matrix of 365 neurons in the larval mushroom body. Rows and columns correspond to neurons ordered using a simulated annealing method for cluster visualization. Pixel intensity corresponds to edge weight on a log scale. Colors highlight within-cluster edges. Neuron ordering is preserved across panels to show the hierarchical organization: with increasing *χ*, larger clusters break into smaller, more tightly-linked clusters. **C.** Cell type composition of clusters. Clusters are ordered by size; clusters consisting of 1 or 2 neurons are not shown. Each hemisphere is separated into two primary clusters, with one containing the input/output neurons and the other containing the sensory projection neurons.

Increasing the resolution by increasing *χ* reveals a nested hierarchy in the hemispheres that follows anatomy and cell type. Higher values of *χ* separate each of the two hemispheres into anterior and posterior halves (Figure 1A,B). These clusters are also composed of distinct cell types. The network we analyzed is composed of 201 Kenyon cells (KC), 48 output neurons (MBON), 30 input neurons (MBIN), 66 projection neurons (PN), and 20 cells of various other types [1]. Distinct populations of KCs have different connection probabilities to the other cell types [1], and our method assigns the input and output neurons to one community and the projection neurons to another, with Kenyon cells split between those two communities based on strength of connectivity (Figure 1C). As *χ* increases, a large community persists in each hemisphere composed of KCs, MBINs, and MBONs, reflecting a repeated tightly-connected microcircuit [1] (Figure 1C, Figure S8). These results highlight the contrast between modularity maximization methods and spectral embedding methods of graph clustering. The latter groups neurons based on common connectivity properties; in the larval mushroom body, the clusters found this way are generally composed of a single cell type [10].

At the coarsest scale, our method thus identifies the clear, bilateral structure of the network. However, it also identifies two less obvious bilateral structures and isolates several very weakly connected Kenyon Cells. At finer resolutions, the structure separates into anatomically meaningful subsets with distinct cell type compositions, grouping sensory projection neurons separately from modulatory input and output neurons. The inferred organization is approximately hierarchical; at successively finer resolution, the communities we find are typically nested (Figure 1B). The agreement between the communities we detected and previously-identified anatomical and cell type organization [1] provides support for our method.

We next turn to the Hemibrain connectome, which is two orders of magnitude larger in size.

### In the Hemibrain, Communities at the Largest Scale Correspond to Anatomically Identified Structures

At the coarsest resolution, *χ* = 0, we identified eight communities in the Hemibrain. Some of these communities correspond to individual anatomical structures in the brain, while others span several brain regions (Figure 2A).

**Figure 2.**
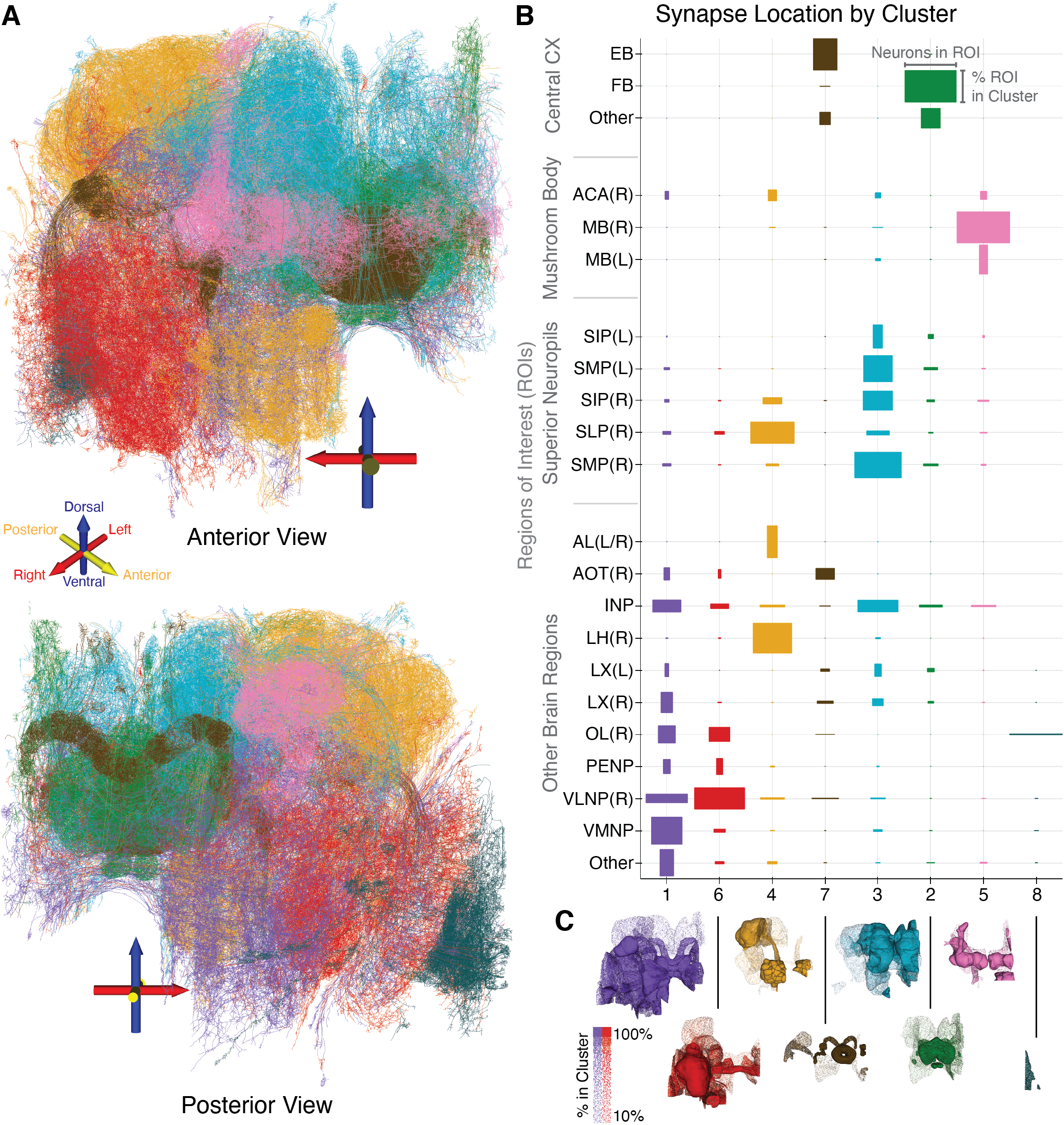
Overview of modularity and anatomy in the Hemibrain. At the coarsest scale modularity, *Q_g_*(0), is maximized by partitioning the Hemibrain network into 8 clusters. **A.** Morphological rendering of neurons color-coded by cluster identity. Shown is a sample of 120 neurons from each cluster (except cluster 8, which consists of only 88 neurons). **B.** Comparison between community structure and anatomy. Brain regions identified by anatomists partition the Hemibrain into disjoint volumes, listed on the y axis. Names are those assigned in the Hamibrain data set, with L/R specifying left and right, respectively. Box height is the fraction of the volume on the y-axis contained in the cluster on the x-axis. The heights in each row sum to 1. The width of a box corresponds to the fraction of neurons in the cluster with synapses in the brain region given on the y axis. Individual neurons may have synapses in many brain regions, so the widths in one column may sum to more than 1. **C.** Volumetric renderings of the brain regions contained in each cluster, with shading indicating only partial containment.

The fly brain is composed of many interconnected neuropils, which have been further partitioned into regions of interest over the last decades by expert anatomists [26]. In Figure 2B, we compare the communities we identified with the previously identified brain regions in the Hemibrain. Box height indicates brain region containment in a cluster, while width indicates neurons with synapses in a brain region. We find that clusters we identified have synapses in restricted subsets of previously-identified regions, and many of those regions are primarily contained in one cluster.

Several anatomical structures in the brain correspond to an entire community identified at this coarsest scale (Figure 2C). For example, the Mushroom Body (MB(L/R)), a structure which plays a role in olfaction and memory [50], forms a single cluster (cluster 5). The fan-shaped Body (FB), which plays a role in spatial navigation and modulation of internal states [23], likewise forms a single cluster (cluster 2). Other structures jointly form a larger cluster, such as the right Lateral Horn (LH(R)), Antennal Lobe (AL(R)), and Superior Lateral Protocerebrum (SLP(R)), which form a cluster with connections to other lateral structures on the right side of the Hemibrain (cluster 4). This is a primary sensory pathway by which olfactory information is transmitted from the antennae to the central brain [41, 53, 27]. The SLP is part of the superior neuropils, which are split into two clusters (clusters 3 and 4). The right superior intermediate and medial protocerebra (SIP(R), SMP(R), respectively) cluster together with their left pairs in cluster 3. This cluster makes bilateral connections to the left and right superior and inferior neuropils (INP). The anterior visual pathway, which transmits polarized light information from the optic lobe to the ellipsoid body [39, 37], forms a single cluster (cluster 7). The remaining ventral neuropils and optic lobe are mostly split between clusters 1 and 6. The lobula plate, a structure in the optic lobe which is not fully contained in the Hemibrain volume, forms a cluster consisting of just 88 cells (cluster 8).

Much of the network topology of the fly brain connectome is closely tied to its spatial topography [25] (see also Figure S9). However, the community structure we find is not simply a reflection of spatial proximity. The anterior and posterior visual pathways (cluster 7 and part of cluster 6), for example, span nearly the whole width of the Hemibrain volume. Likewise, clusters 3, 4, and 5 (making up the superior neuropils and the mushroom body), while spatially colocalized, reflect the distinct roles of their constituent structures in the network: the superior intermediate and medial neuropils (SIP, SMP) form the primary pathway connecting the fan-shaped body to the mushroom body, while the superior lateral protocerebrum (SLP) is part of a network connecting the antennal lobe to the mushroom body.

Thus, at the coarsest level of resolution our algorithm automatically identified well-known anatomical subunits of the fly brain. However, it also placed some anatomical structures with distinct but related functions into the same cluster, suggesting that these structures are tightly linked. We next show that increasing the partitioning resolution can resolve the finer structure of the fly brain, and reveal the hierarchical organization of communities.

### Generalized Modularity Density Reveals the Hierarchical Organization of the Fly Brain

Clustering is a form of dimension reduction for network data that allows us to project a larger network onto a network of clusters and the connections between them. The reduced graph of the clusters we found at the coarsest level, *χ* = 0, reveals high-level organization of the network. In Figure 3A each node represents one of the eight clusters, with the size of a node indicating the number of neurons in the corresponding cluster. The thickness of the edges is proportional to the weighted edge density between clusters (black) and within clusters (gray). Weighted density is how the measure of connection strength is defined for the Generalized Modularity Density, *Q_g_*(*χ*), and we see the within-cluster connections are an order of magnitude stronger than between-cluster connections. Indeed, if we used the same edge scale for both within and and inter-cluster connections, the latter would be nearly invisible.

**Figure 3.**
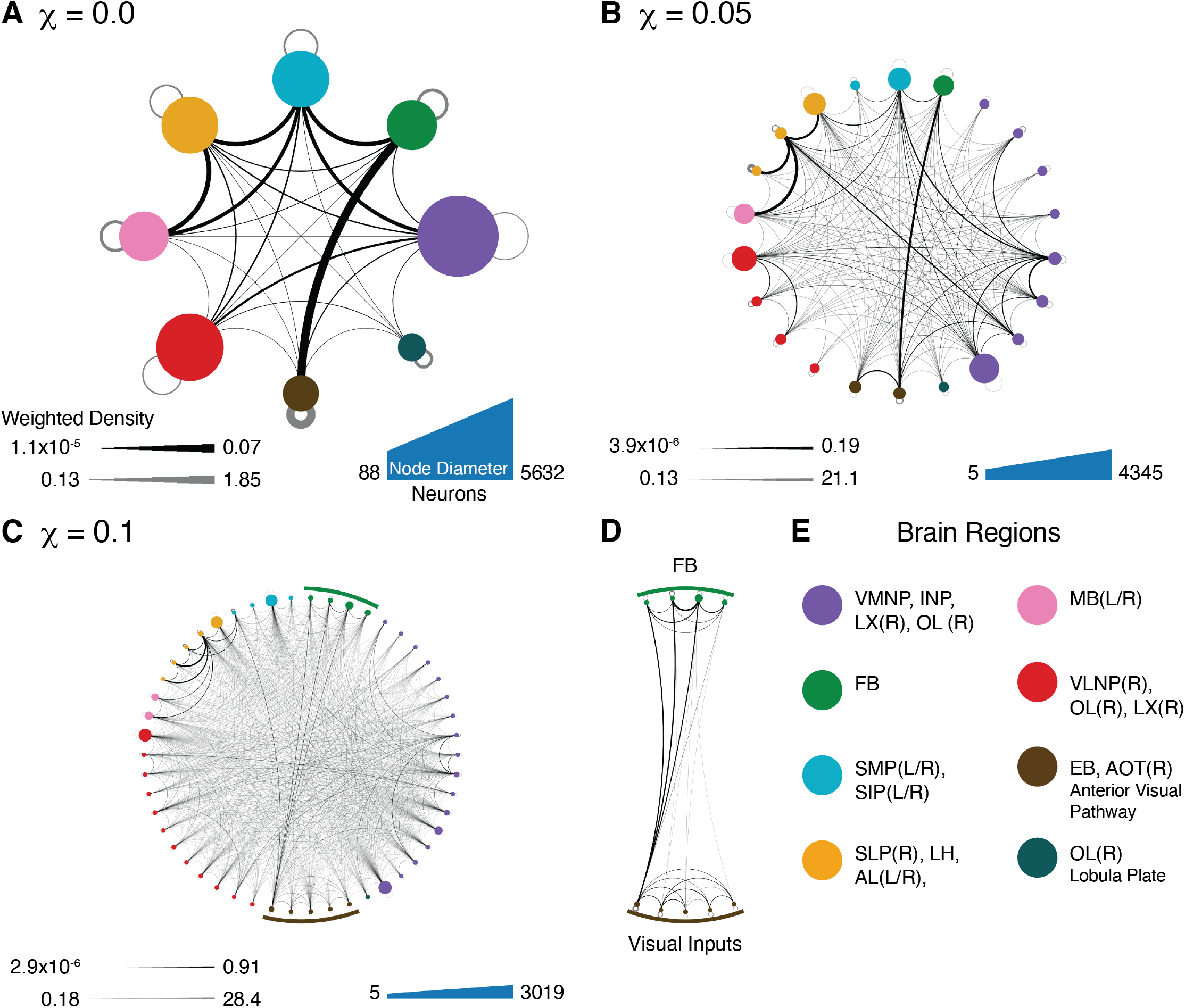
Reduced graphs obtained by representing communities by a single node, and collapsing all edges between nodes in different communities onto a single, weighted edge. Communities with fewer than 5 neurons are not shown. We re-ran the clustering algorithm with *χ* = 0.0, 0.05, 0.1 producing results shown in panels **A**, **B**, **C** respectively. Thickness of edges corresponds to weighted edge density, defined as the total weight of the edges connecting nodes in one cluster to nodes in another, divided by the number of possible edges between the two clusters. The gray border loops represent the connections between nodes within the cluster. The strength of within-community connections is orders of magnitude stronger than between communities. **D.** Clusters making up the fan-shaped body (FB) and anterior visual pathway are isolated to show the organization of the anterior visual pathway – fan-shaped body circuit. **E.** Brain regions represented by the different colors in this figure.

We re-ran our RenEEL community detection algorithm using several values of *χ* to find smaller, more densely-connected clusters. The clusters identified using the classical modularity score, *Q* = *Q_g_*(0), roughly broke into smaller clusters as we increased χ, revealing more specialized sub-clusters. While increasing *χ* generally resulted in a refinement of the partition (i.e. a community *C_i_*(*χ*) at a larger value of *χ* is usually a proper subset of some community *C_j_*(*χ*’) at a smaller value, *χ*’ < *χ*), some clusters appearing at higher values of χ are composed of subnetworks from two or more clusters at lower values of *χ*. However, the vast majority of clusters were broken into smaller subclusters: The majority of clusters identified at resolutions *χ* > 0 were at least 85% contained in one of the 8 clusters found with *χ* = 0 (Figure S14).

Thus, increasing the resolution of the modularity measure allows us to identify sub-clusters that may have important functions in the network. For instance, Figure 3A shows the strong within-community connections in the anterior visual pathway (represented by the thick gray loop on the brown node) and its strong connection to the fan-shaped body (the thick black edge connecting brown and green nodes). The anterior visual pathway splits into two stages in Figure 3B (brown nodes). The first stage consists of the inputs from the optic lobe and is very weakly connected to FB, while the second, more densely-connected cluster, contains the ellipsoid body and retains the strong connection to FB. Increasing *χ* resolves these clusters further, revealing the modular organization of the visual inputs and of the fan-shaped body. The visual inputs are organized into a densely-connected network of clusters which feed into a single densely-connected cluster, which in turn connects strongly to three of the four subclusters of FB (Figure 3C, D). The clusters in FB correspond to distinct anatomical layers, which we discuss further in the next section. We discuss the inputs to the visual pathway further in Clustering Reveals Cell type-specific Wiring Patterns.

Our approach shows that the the fly brain is composed of hierarchically organized communities. The function of many of these communities, and the reason for this structure is not completely understood. However, the inferred structure suggests a functional role for this organization that could be tested in followup experiments, We next take a closer look at the organization of the the fan-shaped body revealed by our method.

### Clustering Automatically Reveals Layering in the fanshaped Body

Having identified the fan-shaped body (FB) as a single community at *χ* = 0, we next investigated the finer structure of this community by increasing *χ*. The FB splits into several sub-clusters, which correspond to previously identified anatomical layers.

At the coarsest resolution FB comprises a single cluster (cluster 2, Figure 2C). This cluster consists of 2391 cells of which 2315 (97%) have synapses in FB; this represents 90% of the 2570 cells in the Hemibrain volume with synapses in FB. This cluster remained coherent at higher values of *χ*: On average new clusters containing any of these initial 2391 cells for *χ*>0 were 95% contained in the original cluster (Figure S14). The sub-clusters we found are arranged from dorsal to ventral, with increasing resolution producing finer layering.

The fan-shaped body is known to have a layered structure [54, 32, 21], with different layers playing different functional roles in the brain [23, 32, 14, 28]. The exact number of reported layers in FB varies, so we compared our results to the 9 layers identified in the Hemibrain data set [45]. Increasing *χ* from 0 to 0.1 split FB into four communities, which roughly correspond to layer 1, layer 2, layers 3 to 6, and layers 7 to 9 (Figure 4A). Increasing *χ* further separated these clusters along the dorsal-ventral axis. At *χ* = 0.5, FB separated into 7 layers, combining the Hemibrain’s layers 4 and 5 and layers 7 and 8 into single communities.

**Figure 4.**
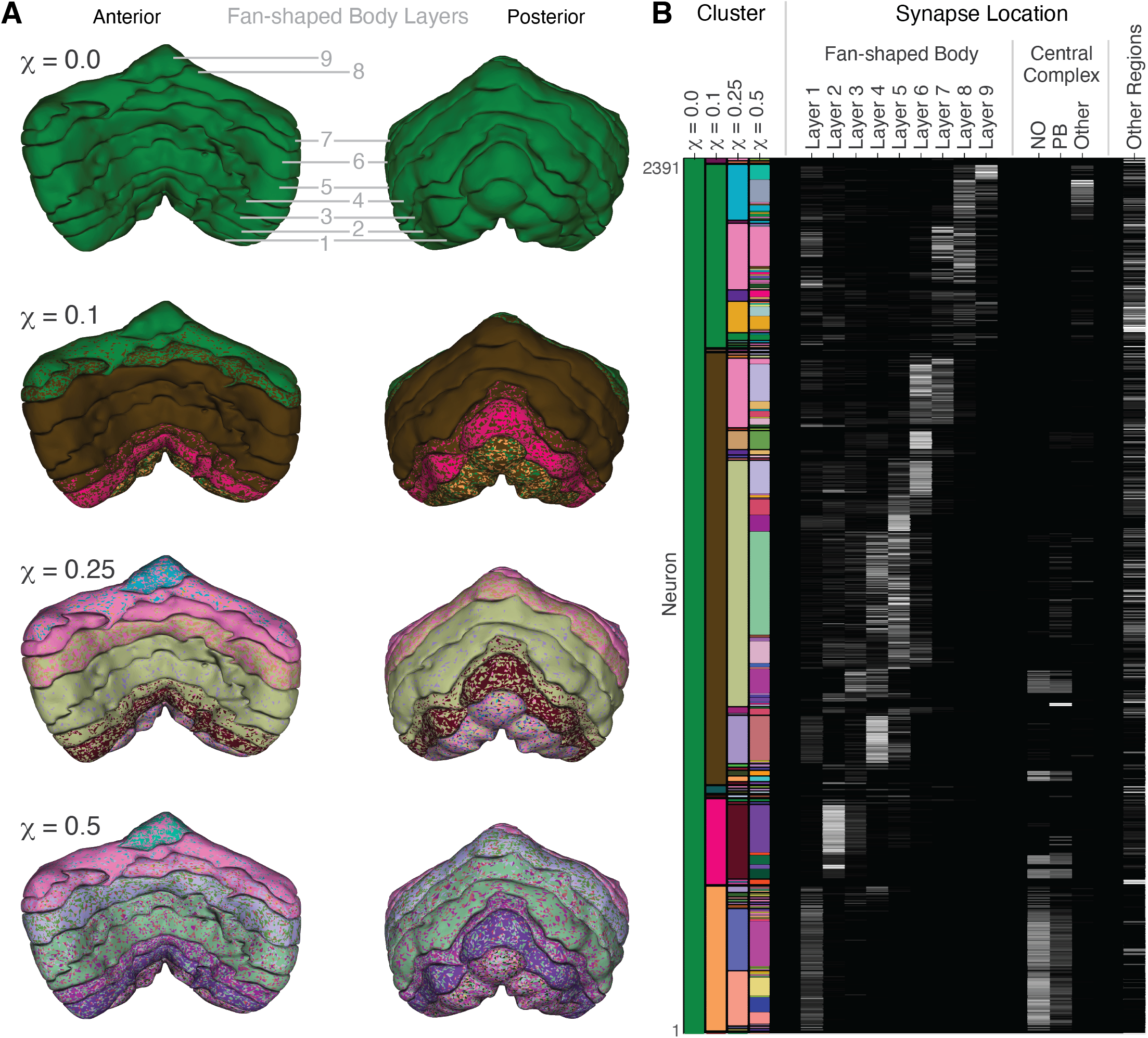
The layered, hierarchical structure of the fan-shaped body (FB) is automatically revealed by varying the tuning parameter, *χ*. **A.** FB is partitioned into nine layers in the Hemibrain data set, rendered here as 3D volumes. Each volume is shaded with color proportional to the fraction of the volume of the corresponding cluster. Left column shows anterior view, right column shows posterior view. Note that at higher resolutions (*χ* > 0.1), most layers are composed of multiple clusters, revealing structure within layers. This structure is reflected in the multicolor texture of each layer. Each of these individual clusters resides primarily in one or two layers, so different layers have different mixes of colors. **B.** Synapse locations of the neurons in cluster 2 (*χ* = 0). Neurons on the y-axis are sorted according to cluster membership at increasing values of χ; clusters are indicated by colors in the left columns labeled with values of *χ*. Repeated colors within one column indicate the same cluster. Most clusters are nested hierarchically, so there are few repeated colors within each column. In the remaining columns, the grayscale intensity shows the fraction of that neuron’s synapses in previously anatomically identified regions listed on the x-axis. For example, the dark pink cluster at *χ* = 0.1 (near the bottom of the figure) resides primarily in layer 2, with projections to the Noduli (NO) and protocerebral bridge (PB).

The strong separation between the dorsal and ventral halves may reflect their different functional roles – the ventral layers play a role in navigation, while the dorsal layers modulate arousal [33, 14, 28]. The parameter χ controls the sensitivity of the clustering to within-cluster density, so our findings show that the anatomically-defined layers are also densely connected as networks. However, the individual layers themselves do not form single network communities. Rather, each layer is composed of a mix of densely-connected networks that reside primarily in that layer.

Thus, our method automatically discovered the layered structure of the FB in an unsupervised way. Our results suggest the layers are hierarchically organized: At the coarsest scale, the FB is split into four layers which are further subdivided at increasing resolution. Some of the clusters we identified relate to the columnar structure of the FB, which we revisit below. We next ask how the structures identified by our algorithm relate to cell types.

### Common Cell Types Form Densely Connected Clusters

Understanding the importance of the identified structures and relationships between them becomes difficult as the number of clusters grows. In order to identify meaningful finer-scale networks, we combined our clustering results with cell type data attached to each node in the Hemibrain.

Most neurons in the Hemibrain (over 90%) were previously assigned cell types based on morphology and brain region connectivity [45]. In contrast, the clusters we found were defined using network connectivity alone, without the use of cell type information. We thus used the partitioning of cells by type in conjunction with clustering to uncover potential cell type-specific wiring principles. Unlikely conjunctions of cell types – such as clusters with a wide variety of cell types, or clusters containing all instances of a given cell type – indicate structures that deserve further examination. A cluster with a wide variety of cell types could represent a functional circuit composed of diverse, but strongly interacting neurons. Alternatively, a very homogeneous cluster could reveal cell type-specific wiring principles. We first discuss the cell type distributions within larger clusters, identifying key functional circuits, and then consider how cells of particular types are partitioned by the clustering algorithm.

To quantify the diversity of cell types that constitute a cluster we used *cluster heterogeneity*, measured using Shannon entropy (see Methods for details). Smaller clusters typically have lower entropy than larger clusters as they can contain fewer cell types (Figure 5A). Increasing *χ* partitions the network into smaller clusters on average, and the average entropy across clusters decreases. However, we found that larger clusters tended to have significantly lower entropy compared to *shuffled* data (Figure 5A, left column), obtained by permuting the cell types between the cells. The largest clusters we found at higher values of *χ* have only a few bits of entropy, so they must be composed of many cells belonging to only a few cell types. This suggests that neuron types that are prevalent in the brain (i.e. types with tens or hundreds of exemplars) are most densely connected with other common cell types, much more so than would be seen in a randomly-labeled network. Therefore, clusters are more uniform in their composition than expected by chance, with some being composed almost completely of one or a few cell types.

**Figure 5.**
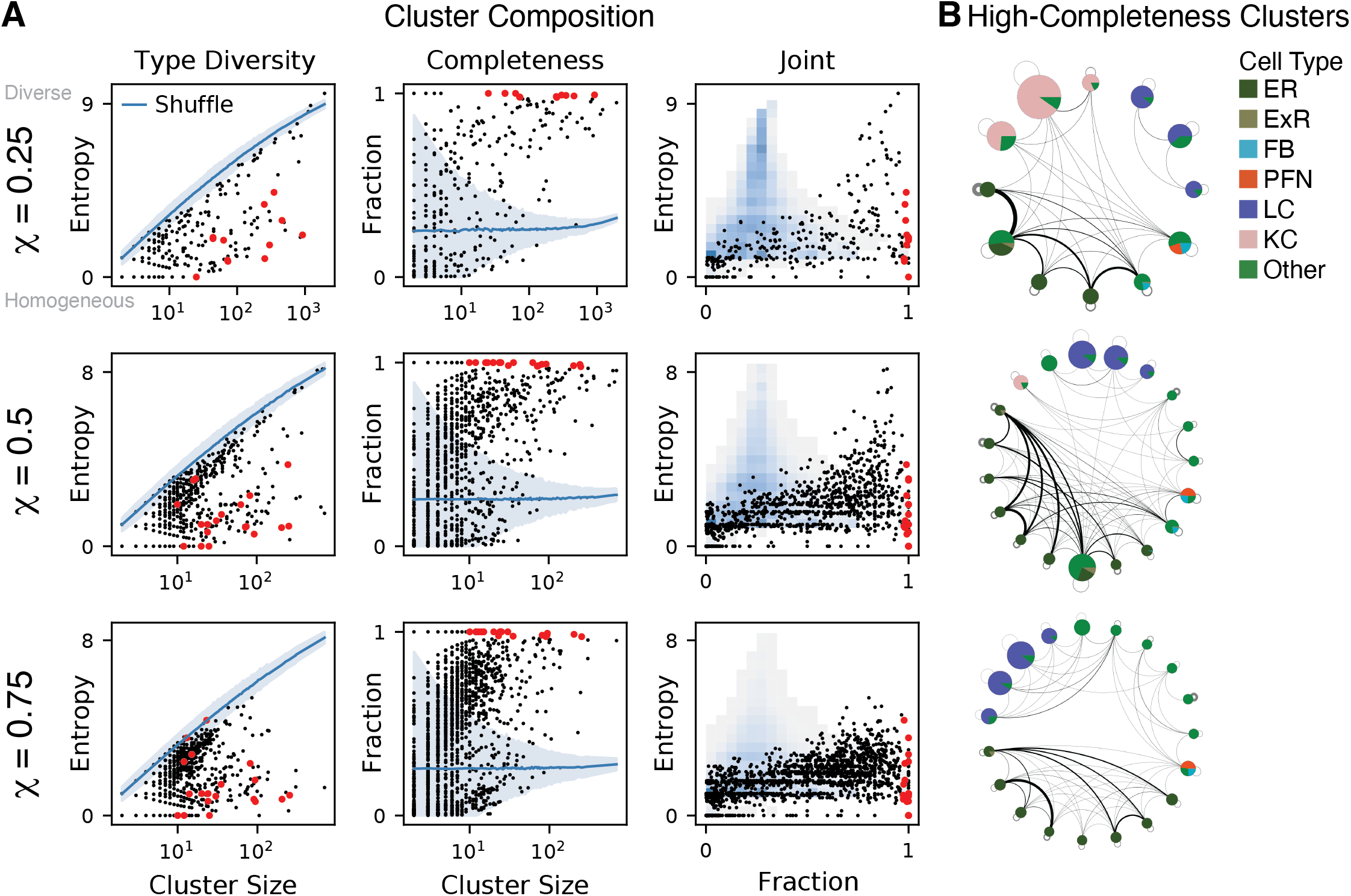
Cell type composition of clusters identified by maximizing *Q_g_*(*χ*). **A.** Heterogeneity and completeness of cell types in clusters. Each row shows clusters identified using a different value of *χ*. The left column shows the heterogeneity (entropy) of cell type distributions within each cluster. Each black dot represents a cluster found using our method, with the corresponding cell type entropy plotted against cluster size. Large red dots indicate clusters shown in networks in panel **B.** The blue line shows the average entropy of clusters in the shuffled data (*n* = 100 shuffles), with the band indicating 3 standard deviations about the mean. The middle column shows cell type completion within each cluster. The right column shows heterogeneity against completeness for the same data; black dots correspond to the actual data, shaded squares show a 2D histogram obtained from shuffled data. **B.** Reduced networks of clusters with at least 97% cell type completion and at least 10 neurons. Node size is proportional to the number of neurons in the cluster. At different resolutions, ER cells and LC cells remain completely clustered (see text).

Cluster heterogeneity does not reveal the degree of homophily among the different cell types, that is, whether common types of cells are most densely connected to other cells of the same type, or whether they tend to share connections with different type cells. To answer this question, we introduced *cell type completeness* of a cluster, defined as the fraction of cells of a given type that belong to a cluster, averaged over the cell types in that cluster (see Methods for details). A cluster with a completeness score of 1 may be composed of multiple cell types, but it contains all cells of those types. For each value of *χ* > 0, we found an average completeness score of 0.4 (the 8 clusters found at *χ* = 0 have an average completeness score of 0.8). Larger clusters tend to have higher completeness scores; excluding clusters with fewer than 10 neurons raises the mean completeness score to the range 0.5-0.7 and clusters with at least 100 neurons have a mean completeness score of 0.8-0.9 for *χ*>0 (Figure 5A, middle column, Figure S12). In other words, certain common cell types form strongly connected communities and these are automatically identified by our clustering method (Figure 5A, middle column).

We found high-completeness clusters distributed throughout the brain, comprising a variety of cell types. The clusters with the highest completeness scores reside primarily in the mushroom body, in the inputs from the visual system, and in the central complex. The two most common cell types in the data set are lobula columnar cells (LC) and Kenyon cells (KC) which combined represent 18.5% of the neurons in the Hemibrain. LCs form the bulk of the portion of the optic lobe in the reconstructed volume of the Hemibrain. There are many subtypes [38, 40], the most prevalent of which are grouped together by our method across multiple resolution scales (Figure 5B). KCs form the bulk of the mushroom body and the majority of the high-completeness clusters in that region. They are densely, but weakly connected [48], and thus do not completely cluster together at higher values of *χ* (Figure 5B).

In the central complex, many of the ER cells which compose the head direction circuit in the ellipsoid body form cliques, all-to-all connections with cells of the same type [44, 45, 15]. These are automatically identified by our method (Figure 5B, dark green). At the coarser resolutions, we find that nearly the entire central complex forms a small number of high-completeness clusters while at the finest scale, the communities we identified are composed of one or two types of ring cells (Figure 5B, bottom; Figure S13).

Cluster heterogeneity and cluster completeness are complementary measures that summarize the cell type composition of clusters. Heterogeneity measures the diversity of cell types within the cluster, while completeness measures how strongly each type is represented. Plotting cluster heterogeneity against completeness organizes clusters into four quadrants allowing us to identify functional networks in an semi-automated fashion (Figure 5A, right column). In the top right (high heterogeneity, high completeness) are clusters which are composed of nearly all cells of multiple types, potentially indicating important functional circuits. Clusters in the bottom right (low heterogeneity, high completeness) are communities composed primarily of a single cell type. In the bottom left (low heterogeneity, low completness) are tightly linked subnetworks of a single cell type, or potentially cell sub-types. In the top left (high heterogeneity, low completeness) is noise – only the shuffled data appears here. Interesting high-completeness clusters are easiest to identify with this framework, as these are the clusters composed of nearly all the cells of one or more types. Clusters in the lower left – low-heterogeneity, low-completeness – require a slightly different analysis, which we turn to next.

### Clustering Reveals Cell type-specific Wiring Patterns

By definition, a high-completeness cluster contains most cells of certain types in the brain. However, the abundance of low-completeness clusters suggests that many cell types form multiple densely-connected sub-clusters which may or may not be homogeneous. This may be a result of fine-scale organization within the cells of one type, such as submodules composed of cells of a single type, or repeated wiring patterns that could be revealed by similarly structured clusters composed of multiple cell types. Our approach can reveal these structures by partitioning cells of a given type into separate clusters with increasing values of *χ*. These separate clusters would have low completeness scores, as they would contain only a fraction of the cells of one type. Moreover, such clusters might only become apparent at high resolution when clusters are smaller and more densely connected. Thus, we sought to identify cell-type-specific wiring patterns by examining the partitioning of cell types into clusters and the dependence of such partitioning on the resolution scale.

Such identification of wiring patterns requires us to quantify the *cluster composition of cell types*. That is, we are interested in finding multiple clusters which share cells of a given type in order to understand the wiring patterns of that cell type. We do so by leveraging the multiresolution nature of our modularity density measure, considering the progressive partitioning found by increasing the control parameter *χ*.

To this end, we considered the homogeneity and completeness of *cell types* rather than clusters (see Methods). For a given cell type, we define *cell type homogeneity* as the fraction of cells of that type which belong to a cluster, averaged over all clusters that contain cells of that type. For low values of *χ*, which partition the network into a few large clusters, most cell types have a homogeneity score of 1. At this scale, most cell types are contained in one of the big clusters. As *χ* increases and clusters break apart, cell type homogeneity typically decreases, as cells of a given type are split among different clusters.

The complementary *cell type completeness* score is the counterpart to cluster completeness, with the role of cell type and cluster identity reversed. That is, the completeness of a cell type is the fraction of a cluster composed of that cell type, averaged over clusters which contain cells of that type. Thus, a cell type with a high completeness score may be partitioned into many clusters, but those clusters are composed primarily of that cell type.

As *χ* varies, the homogeneity and completeness of a cell type vary as well, tracing out a curve (Figure 6A). For most cell types, this curve starts in the upper left corner (high homogeneity, low completeness) and moves down and to the right as *χ* increases. Again, it is helpful to think of cell types in different quadrants of this plot. In the top left, a single cluster contains all cells of that type as well as many other cells. In the top right, cell type and cluster identity are one and the same; cells of this type are most strongly connected to other cells of the same type and form a single cluster. In the bottom right, high homogeneity and low completeness indicates a cell type that has been partitioned into small, homogeneous clusters. A cell type that transitions through these three quadrants as *χ* is increases is one that consistently clusters together at lower resolutions, then breaks up into densely-connected sub-clusters that predominantly contain cells of that type. Computing the area swept out by the curve provides a simple heuristic for identifying cell types that are possibly hierarchically organized in this way. Many of the cell types with the largest swept area values are those that form high-completeness clusters. We therefore focus on the cell types which tend to be part of low-completeness clusters, that is, those types which are partitioned into subsets by the clustering algorithm.

**Figure 6.**
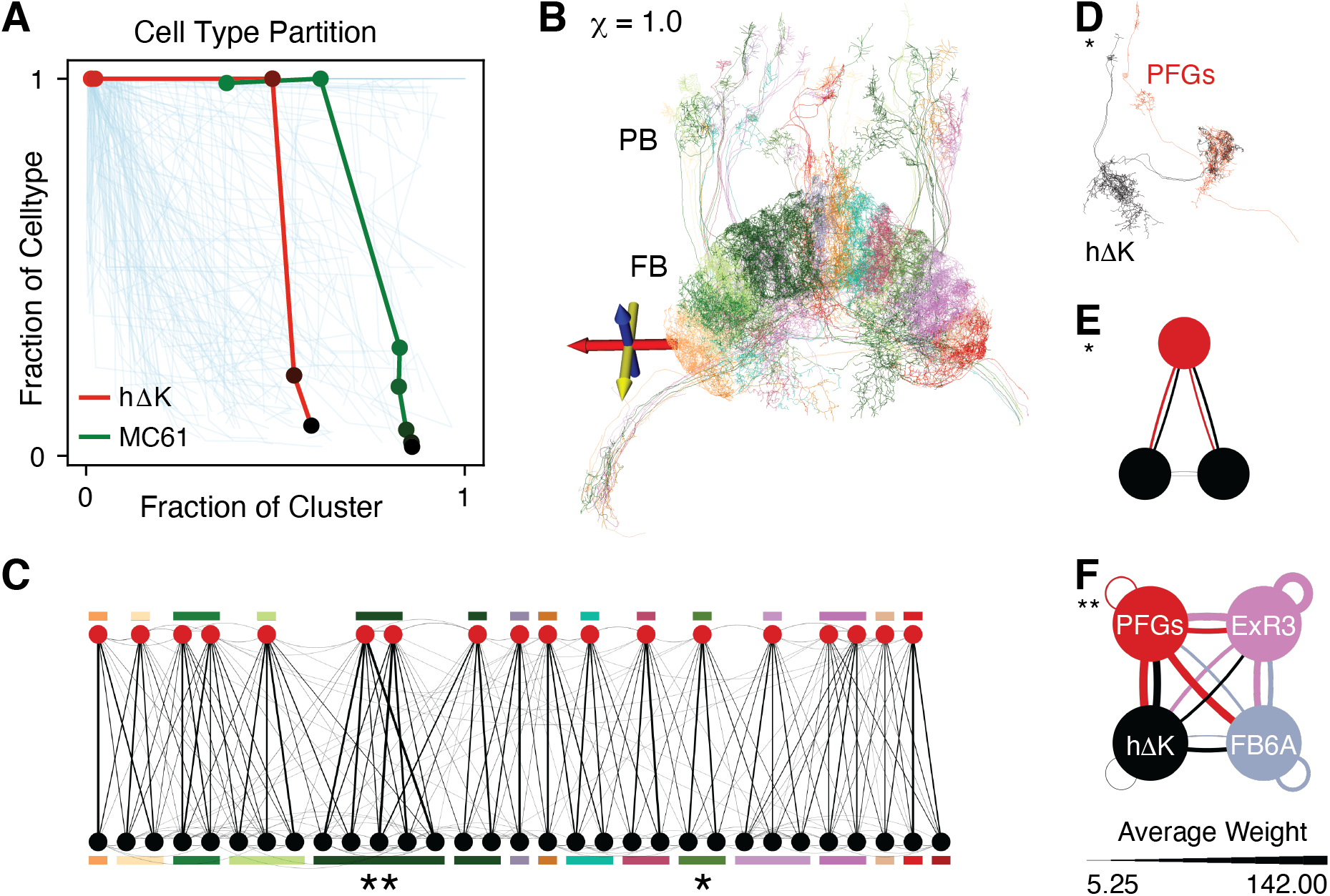
Algorithmically-identified type-specific wiring patterns. **A.** For each cell type, the communities we find partition the cells of that type. This partition is summarized by the completeness and homogeneity scores (see text). As the resolution scale parameter, *χ*, varies, each cell type traces out a curve in the completeness (y axis) vs. homogeneity (x axis) plane. Shown are these curves for all cell types with at least 10 exemplars in the Hemibrain. We highlight two cell types whose curves indicate the possible presence of a repeated type-specific wiring pattern. **B.** Clustering at *χ* = 1.0 reveals the columnar structure of FB. Pictured are all clusters containing hΔK cells, which includes all hΔK neurons and all PFGs neurons. PFGs cells connect the protocerebral bridge (PB) to FB, while hΔK cells are local to the FB. Not shown are five cells which are part of these clusters (a left-right pair of ExR neurons and three FB neurons), but would otherwise obscure the figure. **C.** The undirected network of hΔK neurons and PFGs neurons in these clusters. The network is roughly bipartite, with strong connections between cells of different types, but only sparse and weak connections among cells of the same type. Each cluster is composed of one or two PFGs cells and several hΔK cells. The 18 PFGs cells (top row) are ordered according to their spatial arrangement in FB. **D.** Morphological rendering of the cells in the cluster marked *, colored by cell type. **E.** Directed network representation of cluster marked *, with edges colored by source cell. **F.** The cluster marked ** contains 4 additional neurons, two ExR3 neurons and two FB6A neurons. These neurons form reciprocal connections to all hΔK and PFGs cells, in all clusters. Shown here is the cell type connectivity graph of the indicated cluster. Nodes represent cell types, and edge weights are given by the average connection strength between cells of the two types. Edges are colored according to the source node.

Using this area heuristic, hΔK and PFGs neurons stood out as cell types which potentially partition into type-specific wiring patterns (Figure 6A). The fan-shaped body has a grid-like layout with distinct layers and columns [21, 23, 32]. For higher values of *χ*, the clusters containing the hΔK and PFGs cells tile the middle layers of FB, with each cluster composed of a small number of cells which arborize in two contralateral columns (Figure 6B). PFGs cells connect the protocerebral bridge (PB) and FB; each PFGs cell arborizes in one glomerulus of the PB and one dorsal columnar patch of FB [23]. Each hΔK cell arborizes in two contralateral columns of FB (Figure 6D). Together, these two cell types form a roughly biprartite network with sparse, weak connectivity between cells of the same type (Figure 6C). At the finest resolution scale we examined, corresponding to *χ* =1, Each of these clusters consists of at most two PFGs cells and at most three hΔK cell per PFGs cell. One cluster contains an additional 4 neurons, a pair of ExR neurons and a pair of FB6A neurons, which are strongly, reciprocally connected with all hΔK cells and all PFGs cells (Figure 6E).

Investigating another cell type with a high area score revealed spatial and circuit organization in the inputs to the anterior visual pathway. The small clusters in the anterior visual pathway that emerge at higher values of *χ* (Figure 3D, brown) are composed mostly of MC61 cells, which project visual information directly from the medulla to the central brain [37, 39]. This algorithmically-discovered partition corresponds to a spatial tiling pattern in the small unit of the anterior optic tubercle (Figure 7A). The spatial tiling corresponds to a wiring pattern involving MC61 cells and tubercle-bulb cells (TuBu) (Figure 7B). The clusters that are primarily composed of MC61 cells typically include one or two TuBu cells, which receives input from all MC61 cells in the cluster (Figure 7C). TuBu cells innervate EB ring neurons, conveying visual information to the central complex in a retinotopically organized manner [37, 46]. The striking tiling uncovered by our method, with one tile per TuBu cell, implies a topographic organization of the MC61 dendtritic arbors in the medulla, which is consistent with previous findings [37, 39] Unfortunately, MC61 cells are at the edge of the reconstructed volume and the medulla is not reconstructed, so we cannot describe the relationship between the spatial extent of MC61 cells in the medulla with their cluster identity.

**Figure 7.**
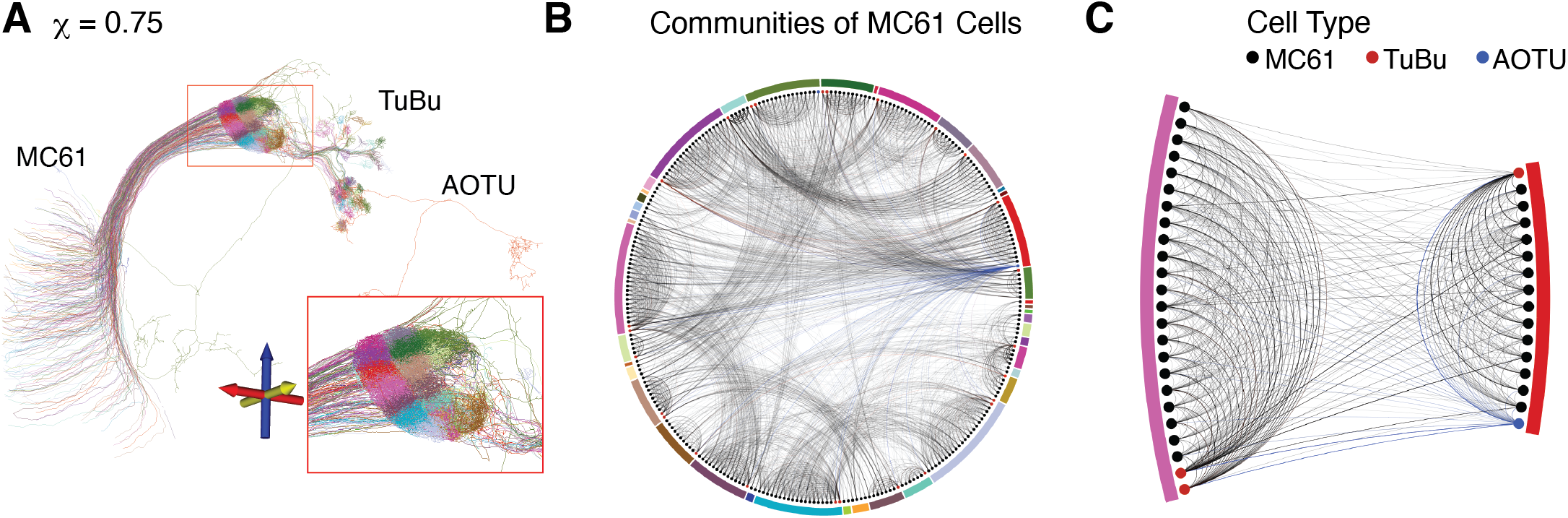
**A.** Clustering at *χ* = 0.75 reveals a distinct spatial tiling pattern in MC61 cells and their outputs, with each tile receiving inputs from spatially congregated input fibers from the medulla. Shown are 293 cells, comprising all clusters which are at least 80% MC61 cells. This includes 258 of 346 MC61 cells (75%) in the Hemibrain data set. **B.** The neurons in **A** form a highly organized network: Each cluster of MC61 cells converges on a common TuBu cell. Two of the clusters are shown in **C**, with edges colored by the presynaptic neuron.

Our method is thus capable of identifying fine, cell-type-specific microcircuits, consisting of as few as three neurons. By comparing the partition of the network induced by cell type annotations to the partitions found by our method across several resolution scales, we can identify cell types which are progressively partitioned into functional modules. The intricate spatial topography revealed by the connectome’s community structure in this way reveals striking fine-scale organization in the fly brain.

## Discussion

We have shown that clustering by maximizing Generalized Modularity Density both recovers known anatomical structures and infers novel organizational principles from connectome data. Our approach allows us to uncover such structure automatically, unsupervised, solely from the network architecture. We also discover cell-type specific wiring patterns when we add node labels added.

Our methods are scalable and can be applied to existing connectomes and to those that will be reconstructed in the future. Our community detection method is fully automated, but biological interpretation is required to fully appreciate the results. Including cell type data with clustering can help generate testable hypotheses about the organization of the network, hypotheses that can be tested using electrophysiology, imaging, genetics [40] and other methods. Thus, our methods are an important tool that can be used to help understand the organization and the function of neural populations and circuits.

There are multiple approaches to finding community structure in complex networks, each identifying communities with different attributes [17]. Maximizing Generalized Modularity Density identifies communities that are more densely connected than a random graph null model [18]. We validated its use for finding connectome structure by analyzing the structure of the Larval Mushroom Body and comparing the results with those found previously with other methods [10]. Our results identified a detailed hierarchical structure of nested communities consisting of heterogeneous cell types in the hemispheres that follows anatomy. The structure we found contrasts with that found using spectral embedding methods, for example, which generally identify single cell-type clusters. Although our method of community detection finds structure that has an appealing, ‘straightforward’ interpretation, it is not necessarily better than other methods. Rather, community structure found by different approaches should be considered complementary.

Brains inherit a degree of hierarchy and modularity during the course of development from neural stem cells [30, 24, 34, 43]. The fly brain, in particular, is composed of clonal units, densely connected populations of cells derived from a single neural stem cell [22, 24]. Since our approach partitions the network into densely-connected clusters, many of the communities we find in the Hemibrain align closely with previously described clonal units [25, 37]. A precise quantitative comparison to the systematic analysis by Ito et al., however, is difficult because we cannot perform a cell-by-cell alignment between their results and ours. Still, there are notable similarities between their Figure 1 [25, p. 645] and our Figure 2: A visual inspection shows that many of the identified clonal units are strictly contained within the clusters we find at the coarsest scale parameter *χ* = 0.

In the Hemibrain, cell body fiber annotations group cells by the location of their cell body on the outer layer of the brain, which correlates with a cell’s clonal origin [45]. At lower resolutions, cell body fibers tend to form subsets of single clusters, while at higher resolutions they tend to be split up (Figure S10). In other species, clonally related neurons may give rise to a large structure such as a cortical column [13], while having a fine-scale organization more similar to a bi- or multi-partite network [5].

By combining our analysis with cell type data, we were able to identify repeated microcircuits composed of as few as three neurons. This relied not only on our ability to overcome the resolution limit problem (e.g. by optimizing *Q_g_*(*χ*) with any particular value of *χ* > 0), but also on our ability to analyze the network at multiple different resolutions. For any fixed partition of the network into communities, a single community of three nodes could potentially be merely random noise. However, the small communities that we highlighted as emerging hierarchically appear to be real biological structures because they have reliable cell-type composition.

It is possible that some of our results are biased by reconstruction errors, and the incompleteness of the data set. For instance, cluster 8 at *χ* = 0 is at the edge of the reconstructed volume, and consists of many neurons which are only partially reconstructed (See Figure 2). We choose to include the cluster in our analysis because it identifies the portion of the lobula plate that is reconstructed in the Hemibrain volume. Similarly, other clusters composed of cells that are only partially contained in the Hemibrain data set could be affected by the incompleteness of the data. In the Hemibrain, automated synapse detection has an average precision of 0.8 and average recall of 0.8, though accuracy varies between brain regions [45]. In order to determine if our results are robust with respect to missing or misidentified synapses, we perturbed a comparable fraction of edges in numerical experiments, and observed no large changes in the identified communities, especially on the larger scale (Figure S15). However, if reconstruction errors are systematic they could change the finer structures detected using community detection methods. Further analysis could be done to estimate the impact of such errors, perhaps by perturbing the synapses in a way mimicking the errors known to occur in network reconstruction [52].

To help us identify communities of interest, we have made use of cell type data in addition to the architecture of the connectome. However, methods that directly incorporate such information could be used to identify network community structure. For instance, spatial information about the cells could be included to preferentially detect clustering between proximal or distal cells. Moreover, we could look at different modes of organization: A core–periphery analysis could identify cores of strongly interconnected cells, and peripheries of cells connected to the core [42]. Similarly, we could search for anti-communities of cells that are connected weakly to each other, but strongly to those in other anti-communities [6]. An analogous bipartite or multipartite method that first distinguishes cells based on their type and then looks for structure within sets of cells of the same type would allow the direct inclusion of cell type information in the clustering, but would require the development of new generalized modularity measures and algorithmic methods.

Neurons do not interact only via synaptic connections, but can influence each other through gap junctions, expression of neuromodulators, and possibly ephaptic and other types of interactions. Moreover, in mammals glia can play an important role in neuronal interactions. Currently available connectomes are based only on synapses between neurons, and thus offer only a partial picture of the interactions between neurons and subgroups. We view our results, and those of similar approaches, as an initial step in characterizing the structure of interactions in the brain, providing a picture that will both change and become more detailed as more complete data becomes available.

## Methods

### Data sets Analyzed

The larval mushroom body data set is a dense reconstruction of all neurons in the mushroom body of a larval fruit fly [1]. The network we studied consisted of 365 neurons, selected for having at least one synapse in the synapse table published as part of the supplemental data of [1]. From the synapse table we constructed a weighted, directed graph with neurons as nodes and edge weights defined by total number of synapses between neurons. For input to the community-detection algorithm, we combined antiparallel edges by summing their edge weights to obtain an undirected graph.

The Hemibrain data set is a dense reconstruction of roughly one half of the central brain of an adult fruit fly, *Drosophila melanogaster* [45]. The network we analyzed was based on version 1.1 of the data set and consists of 21,733 nodes and 2,872,500 undirected, weighted edges. As for the larval data set, the undirected edge weights in the network were the total number of synapses between each pair of neurons, in either direction.

### Detecting Communities with Generalized Modularity Density Maximization

We have identified communities of nodes with clusters *C*_1_,*C*_2_, … in a partition of the nodes *C* = {*C_1_*, *C_2_*, …} that maximizes Q_g_, the Generalized Modularity Density measure [19]:

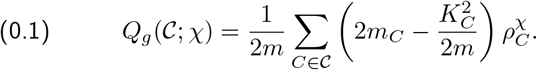

Here *m* is the sum of the weights of all links in the network, *m_C_* is the sum of weights of all links between nodes in community *C*, *K_C_* is the weight-degree sum of nodes in *C* (the sum of the weights of the links connected to each node in *C*), and *ρ_C_* is the relative density of connections in C,

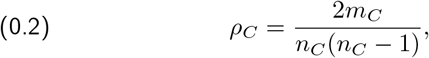

where *n_C_* is the number of nodes in *C*. The exponent *χ* is a tunable control parameter. We consider *χ* ≥ 0. At *χ* = 0, *Q_g_* equals the classical modularity score [36]. Modularity is the difference of fraction of the network’s links that are within communities of the partition C minus what that fraction would be on average if the links were randomly shuffled. For *χ* > 0, a powerlaw function of the relative density of each cluster *C* weights the cluster’s contribution to Modularity in the *Q_g_* measure.

For networks of the size and density we consider here, Modularity (*χ* = 0) is typically maximized by partitions that consist of a relatively small number of large clusters. In fact, it can be difficult to resolve clusters that are small by maximizing Modularity, even if those clusters are very well connected. This well-known resolution limit problem with maximizing Modularity [16, 49] is mitigated by maximizing Generalized Modularity Density at positive values of *χ* [19]. As *χ* is increased the clusters in partitions that maximize Generalized Modularity Density tend to subdivide into smaller more tightly-linked clusters [18].

### RenEEL Algorithm for Maximizing Generalized Modularity Density

Finding the partition that maximizes Generalized Modularity Density can be a challenging and computationally expensive problem, especially for larger and denser networks. An exact solution for an arbitrary network has an NP-Complete computational complexity [3]. Thus, it is necessary to use an approximate method that is fast, with polynomial time complexity, but still gives accurate results. We used an algorithm that has been shown to give accurate results for networks as large and dense as we consider in this paper [18]. This algorithm is based on Reduced network Extremal Ensemble Learning (RenEEL), which employs a machine learning paradigm for graph partitioning, Extremal Ensemble Learning (EEL). ELL evolves an ensemble of partitions toward consensus by replacing the “worst” partition with a new one. RenEEL efficiently generates the new partition by expending effort only where there is disagreement within the ensemble’s existing partitions. The speed and accuracy of our algorithm enables analyses of networks with tens of thousands of nodes, such as ours, that previously were not possible.

The RenEEL scheme for community detection first uses a very fast base algorithm to create an ensemble of partitions that try to maximize a modularity measure. Then an iterative learning process is used to update the ensemble. A reduced network is formed by combining the groups of nodes that the ensemble partitions agree should be clustered together into single nodes. The base algorithm is then used to partition the reduced network. The ensemble is updated either by using the new partition to replace the ensemble partition with the lowest modularity or by reducing the size of the ensemble if the new partition matches one already in the ensemble or has a lower modularity than any in the ensemble. The learning process continues until only one partition remains in the ensemble and, thus, consensus is reached on what partition maximizes modularity. In our implementation of the RenEEL scheme, we used a randomized greedy algorithm [11, 35] for the base algorithm and typically started with 100 partitions in the ensemble.

We tested the robustness of the clustering results by repeating our analysis multiple times and found that consistently clustered pairs of neurons are orders of magnitude more common than inconsistently clustered pairs in the Hemibrain network (Figure S11).

### Information Measures

Clusters define a partition 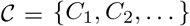 and cell types define a partition 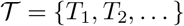 of the nodes in the network. By comparing these partitions we are able to identify potentially biologically relevant networks.

Cluster heterogeneity quantifies the variety of cell types within a cluster, and is measured using Shannon entropy. We defined the heterogeneity of a cluster *C* as

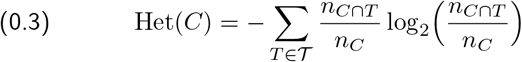

where *n_C_* is the number of cells in cluster *C* and *n_C⋂T_* is the number of cells of type *T* in cluster *C*, and the sum is taken over cell types *T*. If a cluster consists entirely of cells of one type, the cell type distribution has zero bits of entropy, while a cluster composed of equal numbers of cells of n different types has log_2_(*n*) bits of entropy, the maximum for a cluster composed of n types.

Cluster completeness is the fraction of cells of a given type that belong to a single cluster. This is the fraction of a cell type present in a cluster averaged across the cell types within that cluster, defined by

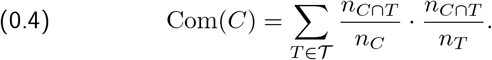

Here *n_C_* and *n_C⋂T_* are the same as in Eq. (0.3), and *n_T_* is the number of cells of type T in the whole network. A completeness score close to 1 means that, among cell types represented in a cluster, nearly all of the cells of those types belong to that cluster.

We defined analogous measures for cell types. Switching the role of cluster and cell type in Eq. (0.4), we defined the completeness of a cell type as

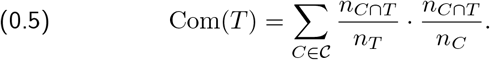

Notice the sum is now over clusters *C*. For cell type, a completeness score near 1 means that clusters which contain cells of type *T* are almost entirely composed of cells of type *T*.

Cell type homogeneity is a measure of the partitioning of cells of a given type. That is, if all cells of a given type belong to one cluster, we say that cell type is very homogeneous. We defined cell type homogeneity as

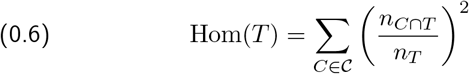

This is the fraction of the cell type contained in a cluster, averaged over clusters which contain cells of that type. A score of 1 means all cells of type *T* are contained in one cluster, with lower scores indicating that cells of type *T* are partitioned into multiple subsets by the clusters.

### Visualization of clusters

To better visualize and compare our clustering results, we used a simulated annealing method to arrange cells in the adjacency matrix plots. In Figure 1B, the clustering results of three different values of parameter *χ* are visualized to show the evolution of clustering as the parameter increases, and the spontaneous emergence of the hierarchical structure. Specifically, for visualization we ordered the nodes to minimize a function *H* of the Euclidean distance *d_ij_* between matrix elements (*i,j*) and the closest point on the diagonal (under periodic boundary conditions). This function takes the form

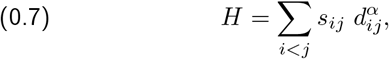

where *S_ij_* is the link weight between node *i* and *j*. Note that we analyzed undirected networks, so the sum is over the elements above the diagonal. The parameter *α* controls the non-linear dependence of H on *d_ij_*. Figure 1B is generated by setting *α* = 3.

For the order we used, we started with the partition at *χ* = 0.5, and used simulated annealing to swap pairs of clusters and pairs of nodes within clusters until the order of nodes and clusters minimized the cost function. Fixing this partition, we then repeated this process with the partition at *χ* = 0.25, now only swapping clusters from the finer partition (*χ* = 0.5) within the clusters defined by the coarser partition (*χ* = 0.25). This process was then repeated for the partitions at *χ* = 0.0 and *χ* = 0.25. The final order thus obtained was used for plotting all three plots in Figure 1B.

### Reproducible research with dynamic figures

The python code and data to generate these figures is available at https://github.com/josiclab/flybrain-clustering. Many of the figures in this paper are simplified to be printer-friendly, and, necessarily, static. They are available online at https://josiclab.github.io/flybrain-clustering/ rendered as interactive plots using javascript so zooming, panning, and mouse-over information is available.

## Acknowledgements

The authors thank Brad Hulse, Romain Franconville, and Fabrizio Gabbiani for helpful discussions.

ABK was supported by a training fellowship from the Gulf Coast Consortia, on the NLM Training Program in Biomedical Informatics & Data Science (T15LM007093). JG and KEB were supported by NSF grant IOS-1546858. ABK, XP, and KJ were supported by NSF NeuroNex grant 1707400. XP was supported in part by the Intelligence Advanced Research Projects Activity (IARPA) via Department of Interior/Interior Business Center (DoI/IBC) contract number D16PC00003. The U.S. Government is authorized to reproduce and distribute reprints for Governmental purposes notwithstanding any copyright annotation thereon. Disclaimer: the views and conclusions contained herein are those of the authors and should not be interpreted as necessarily representing the official policies or endorsements, either expressed or implied, of IARPA, DoI/IBC, or the U.S. Government.

## Author Contributions

ABK analyzed data, produced figures and wrote code. JG and KEB contributed clustering software and analyzed data. KEB, KJ, and XP helped design the study and suggested analyses and visualizations. All authors contributed to writing the manuscript.

## Competing Interests

The authors declare no competing interests.

## Supplementary Materials

**Figure S8.**
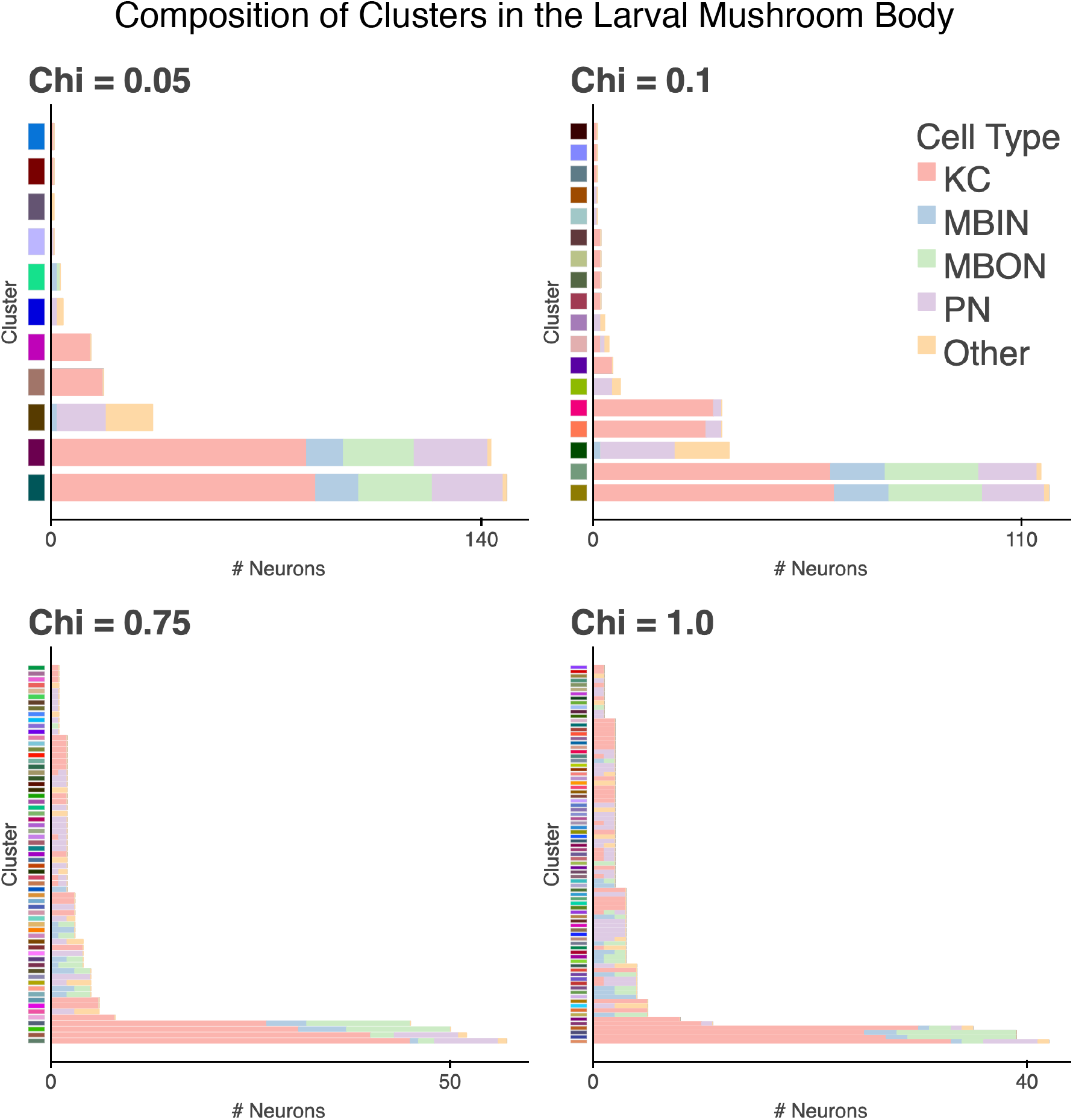
The two hemispheres of the larval mushroom body emerge as the two largest clusters at lower values of *χ*. At higher values of *χ* each hemisphere splits into two lobes with different cell type compositions.

**Figure S9.**
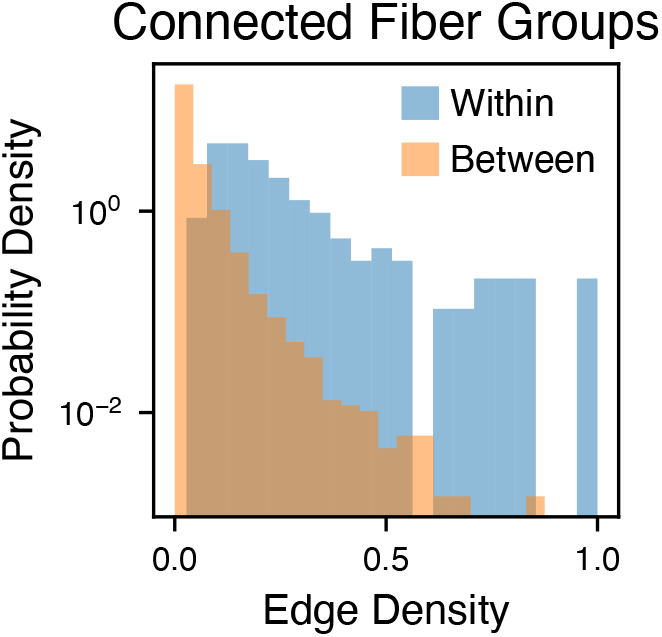
Distribution of edge densities between and among connected cell body fiber (CBF) groups. CBFs are grouped into eight octants (anterior/posterior, ventral/dorsal, lateral/medial), then further divided into groups based on clonal origin [45]. There are a total of 192 CBF groups annotated in the Hemibrain data set. Edge density between groups is defined as the number of edges connecting nodes in one group to nodes in the other, divided by the product of the sizes of the two groups. Within-group edge density is defined as the number of edges connecting two nodes within the group, divided by *n_G_*(*n_G_* — 1)/2, where *n_G_* is the number of cells in the group. Edge density is computed only between connected groups; only 85% of possible connections between CBF groups are present in the brain. The higher edge density within-CBF group compared to between-CBF group reflects the spatial organization of the hemibrain network, that is, the network topology reflects spatial topography.

**Figure S10.**
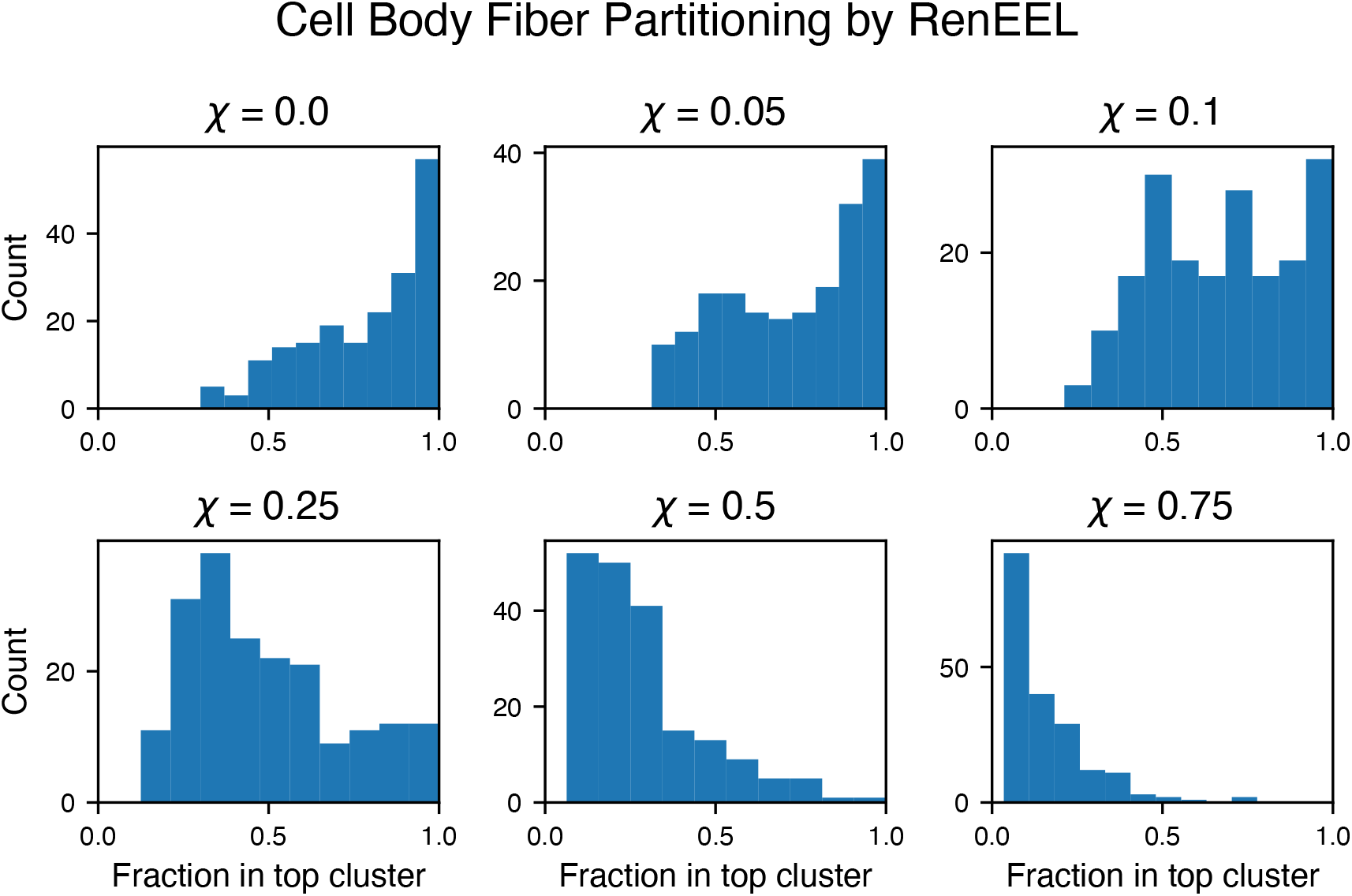
Cell body fiber (CBF) groups partition the neurons in the fly brain based on cell body location and clonal origin. For each cell body fiber (CBF), we computed what fraction of that CBF belonged to each cluster; shown here is the distribution of the top fraction for each CBF. At lower resolutions, RenEEL clusters most neurons in one cell body fiber together, while at higher resolutions, these groups are split among several clusters.

**Figure S11.**
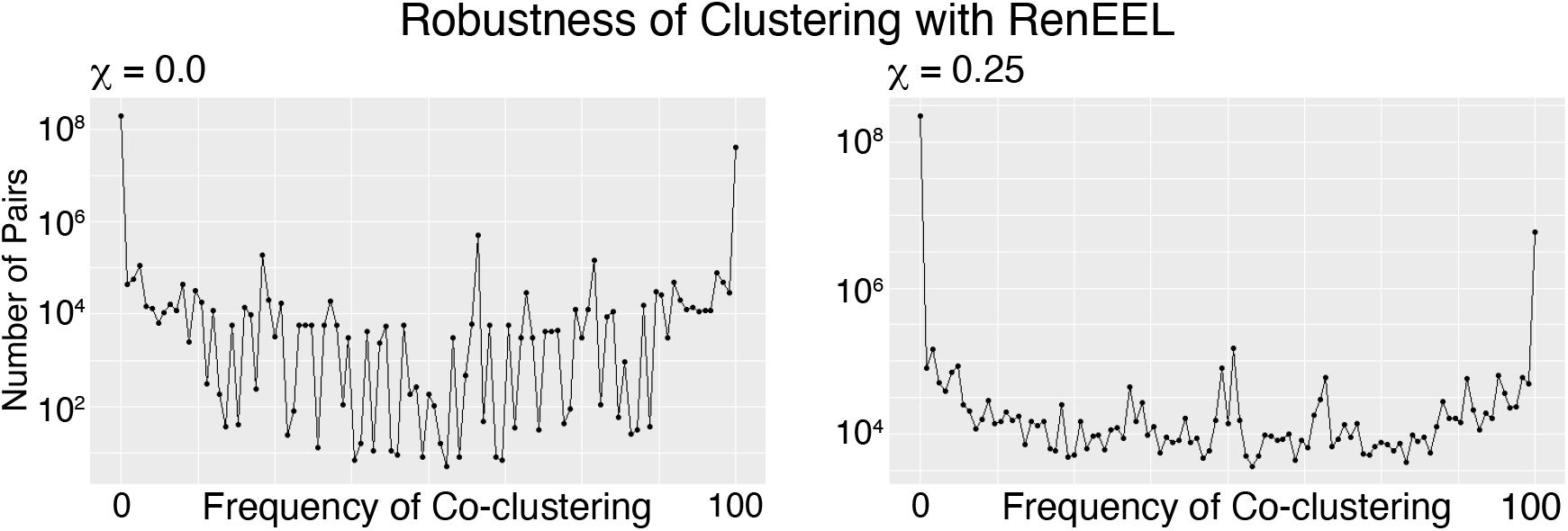
Robustness of clustering. RenEEL was run with a different random seed 100 times; plotted is frequency with which each pair of neurons appears in the same cluster. Pairs of neurons which are consistently clustered have a frequency of 0 or 100. Consistently clustered pairs of neurons are orders of magnitude more common than inconsistently clustered pairs.

**Figure S12.**
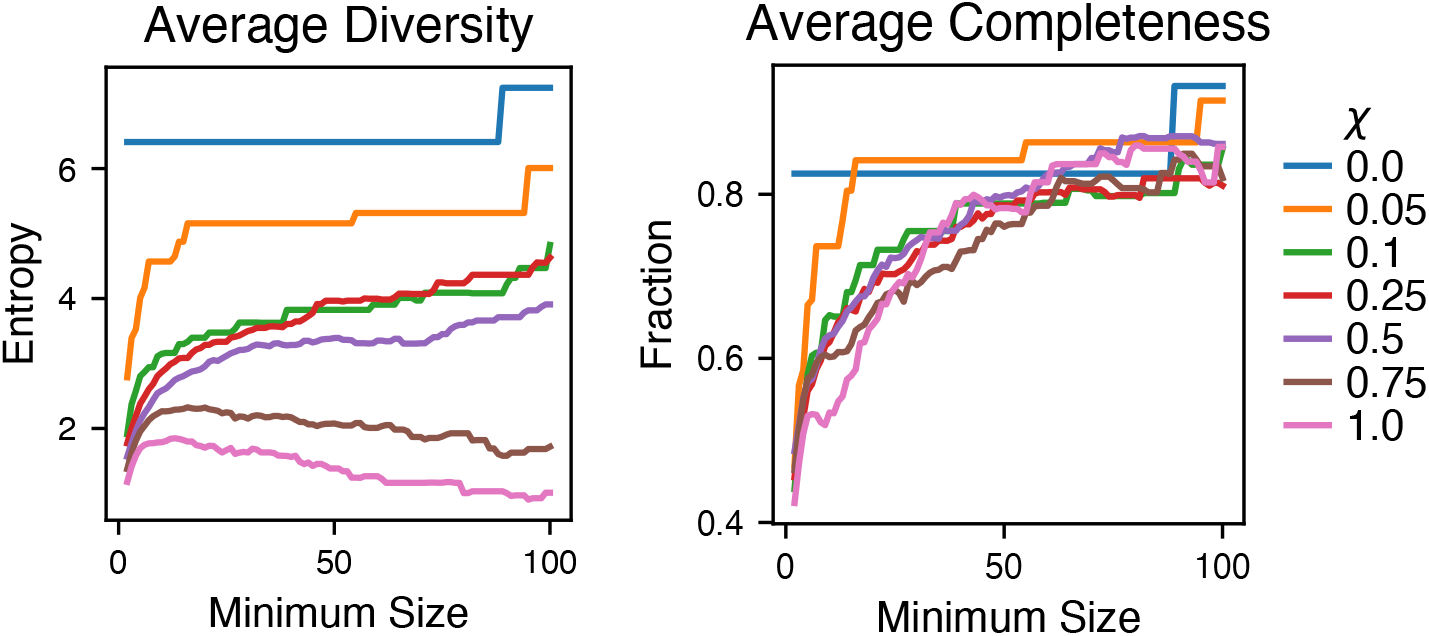
Cell type heterogeneity and completeness (see Methods), averaged across clusters. Shown here is the dependence of this average on the minimum size of clusters included. Cluster heterogeneity decreases as *χ* increases, which reflects the smaller cluster size overall. Cluster completeness remains fairly constant regardless of *χ*. Excluding small clusters raises the average completeness.

**Figure S13.**
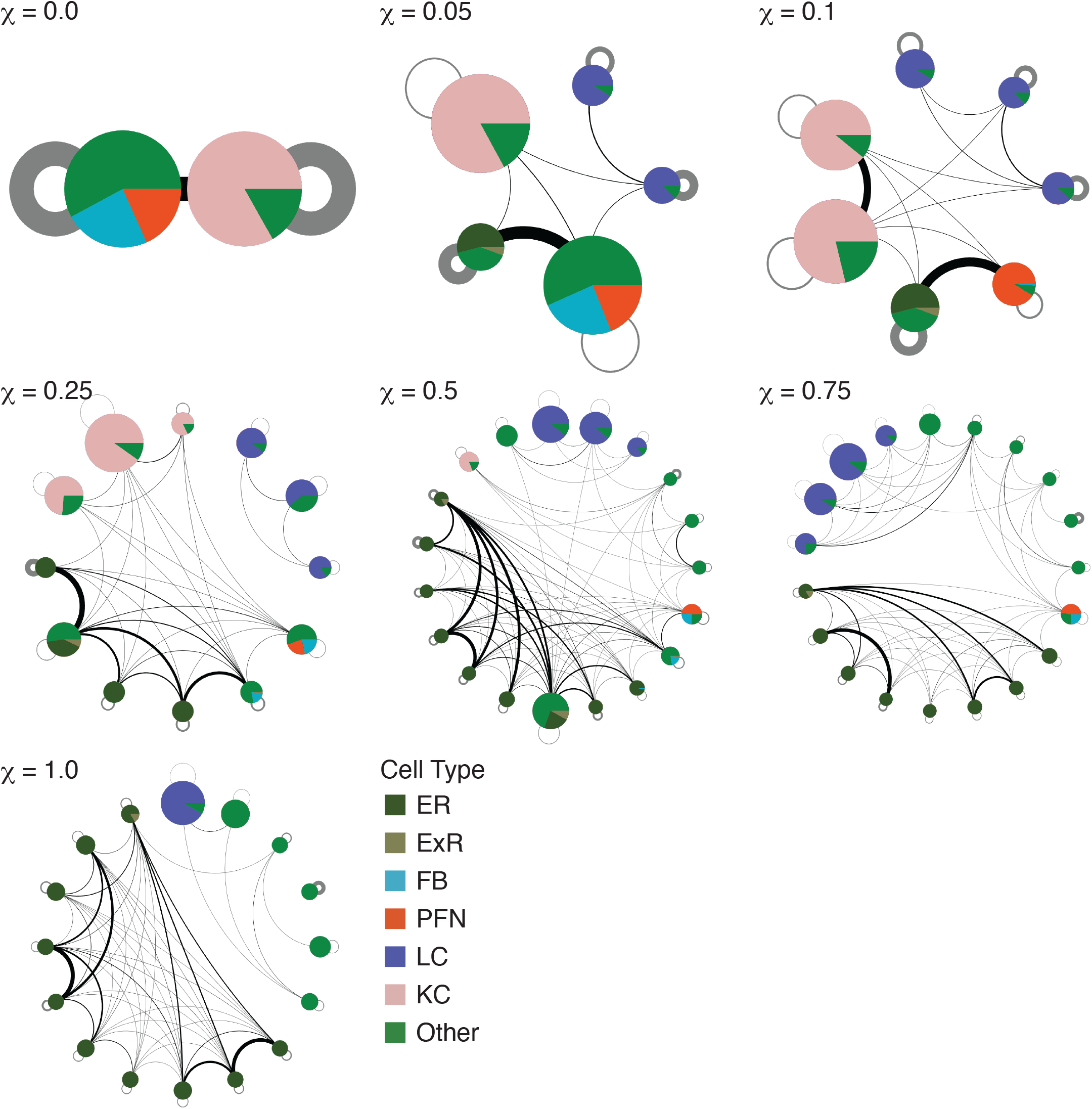
High-completeness clusters for all values of *χ*. Shown are reduced networks for all clusters of at least 10 neurons with a cell type completeness score of at least 0.97.

**Figure S14.**
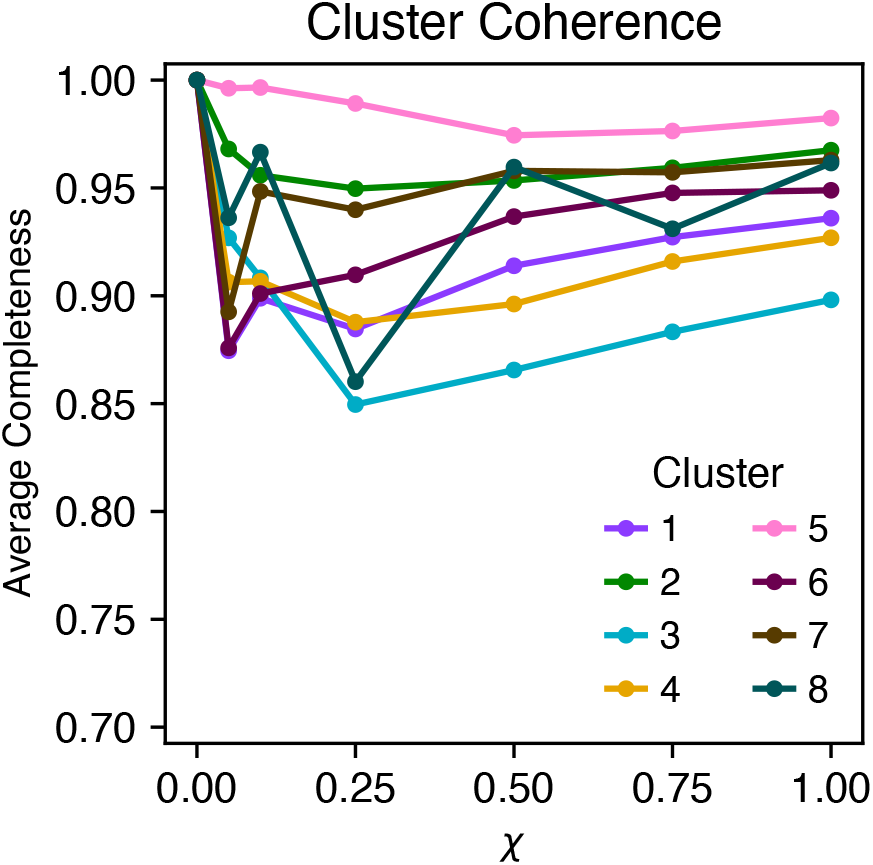
The coherence of each of the 8 clusters found by optimizing *Q_g_* with *χ* = 0. Coherence of one of the eight clusters *C* is defined as the fraction of clusters *C*’ found with *χ*’ > 0 contained in *C*, averaged over the the *C*, which have nonempty intersection with *C*. If the clustering found by increasing *χ* was strictly hierarchical, all values would be 1.

**Figure S15.**
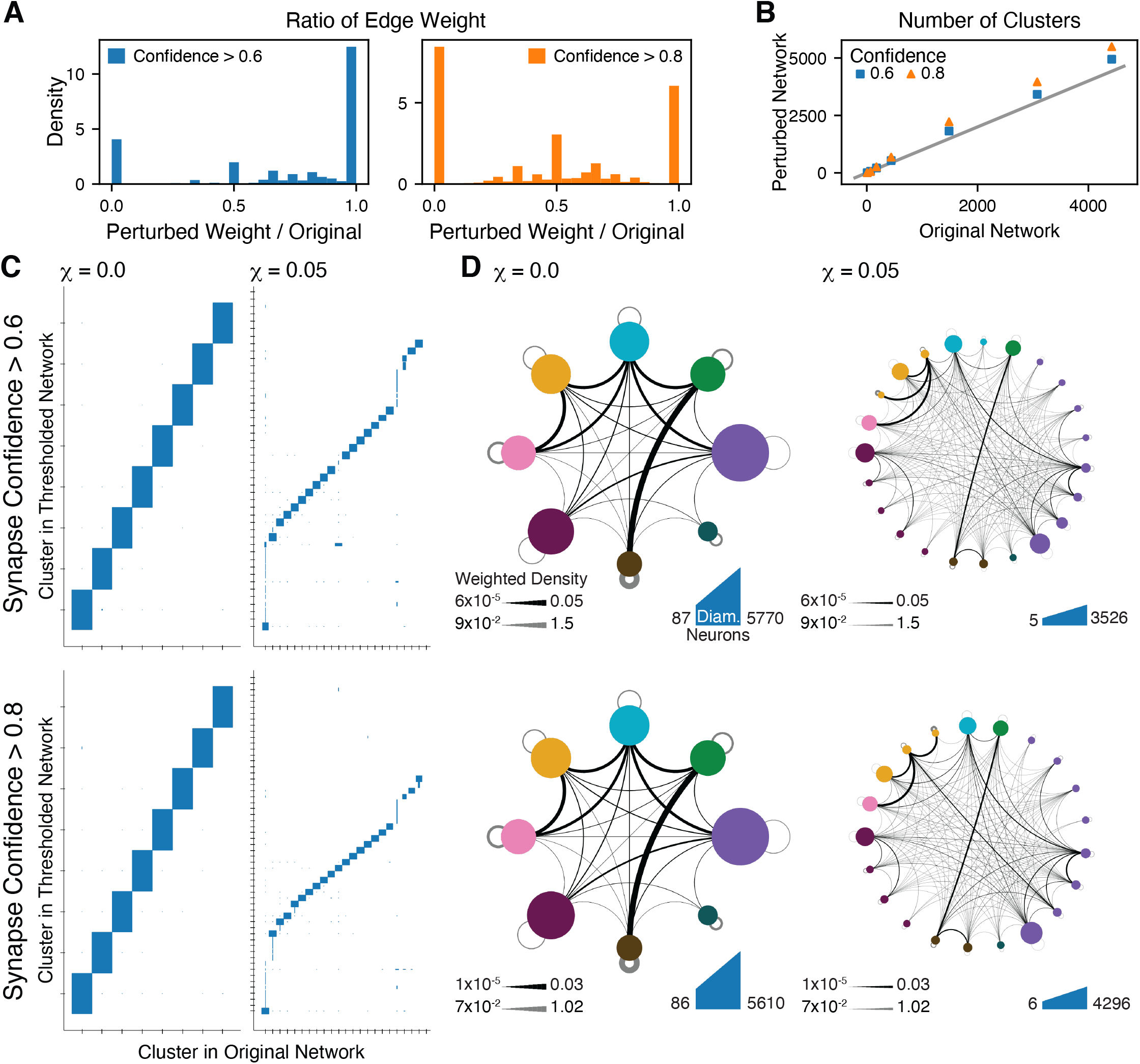
We perturbed the network to investigate the possible effects of reconstruction errors on our results. Community identification at the coarse scale did not significantly change when low-confidence synapses were dropped from the network. **A.** Synapses in the Hemibrain each have a confidence score, indicating the confidence of the machine learning algorithm which automatically identified them. We constructed perturbed networks by omitting individual synapses whose confidence fell below a threshold, then recomputing all edge weights. Each edge in the perturbed network had a weight which was a fraction of its original weight; shown here is the distribution of these weight ratios. This perturbation resulted in overall weaker edges, with higher thresholds also severing more edges (counted in the bin at 0.0). **B.** The number of communities found by our method on the perturbed networks compared to the number found in the original network. Gray line shows equality. The number of clusters found increases with the resolution scale. At higher resolution scales, as the perturbed graph became weakly connected, more clusters were found. **C.** Comparison of the clusters found in the original network with those found in the perturbed network. Each box represents the neurons assigned to the cluster given on the x-xis in the original network and the cluster given on the y-axis in the perturbed network. Box width represents the fraction of the cluster in the original network; box height represents the fraction of the cluster in the new network. Clusters from the original network consisting of fewer than 5 neurons are not shown. At the lower values of χ presented here, the clusters found in the perturbed network are usually wholly contained in a cluster in the original network, shown by the full height of the boxes. **D.** The inter-community connectivity is also fairly similar in the perturbed networks.

## Notes

https://github.com/josiclab/flybrain-clustering

### Competing Interest Statement

The authors have declared no competing interest.

